## Supplementary Tables S1, S2 and Figures S1 - S10 for "Cas9 is mostly orthogonal to human systems of DNA break sensing and repair"

[*]Correspondence should be addressed to OL or DZ

**S1 Table. Oligonucleotides used in this study.**

| Name | Sequence, 5′→3′ |
|---|---|
| ***Cas9 activity assay*** | |
| DNA1 | CTGATAACTCAATTTGTAAAAAATGGTACTGAGCA |
| DNA2 | TGCTCAGTACCATTTTTTACAAATTGAGTTATCAG |
| ***pLK1 plasmid construction*** | |
| pLK1.TOP | TCGAGATAACTCAATTTGTAAAAAATGGTAG |
| pLK1.BTM | AATTCTACCATTTTTTACAAATTGAGTTATC |
| ***Guide RNA cloning into pX458*** | |
| pX458.H4.TOP | CACCTGTCTGGGGACACGTCTCCA |
| pX458.H4.BTM | AAACTGGAGACGTGTCCCCAGACA |
| pX458.H6.TOP | CACCGAATGAAAATGCGGTTCTTG |
| pX458.H6.BTM | AAACCAAGAACCGCATTTTCATTC |
| pX458.H9.TOP | CACCCCGTCACTGAGACAGTGCGC |
| pX458.H9.BTM | AAACGCGCACTGTCTCAGTGACGG |
| ***TIDE analysis*** | |
| TIDE.H4.FWD | TGGCAGGGCTGGTCTTTCTCTGGCA |
| TIDE.H4.REV | AGTCCCGAGCCAAAGCCGAGTGACA |
| TIDE.H6.FWD | TTCTCCTTTCAGATATGGCTGG |
| TIDE.H6.REV | AAGAGATGAATGTATGGGTATGGCT |
| TIDE.H9.FWD | GCTGCCTCTGTTCTTCACCT |
| TIDE.H9.REV | GGTTAAAATGTCACCAGGGTCC |

**S2 Table. PCR protocol for genomic targets amplification.**

| Temperature, °C | Time | Number of cycles |
|---|---|---|
| 95 | 3 min | 1 |
| 95 | 30 s | 35 |
| 72 (H4)<br>62 (H6, H9) | 30 s | |
| 72 | 30 s | |
| 72 | 5 min | 1 |
| 4 | ∞ | 1 |

5′-CTG**ATAACTCAATTTGTAAAAAA**TGGTACTGAGCA-3′ strand 1
3′-GAC**TATTGAGTTAAACATTTTTT**ACCATGACTCGT-5′ strand 2

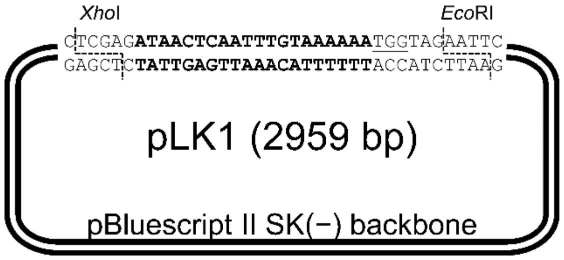


**S1 Fig**. **Substrates used in the Cas9 activity assays.**

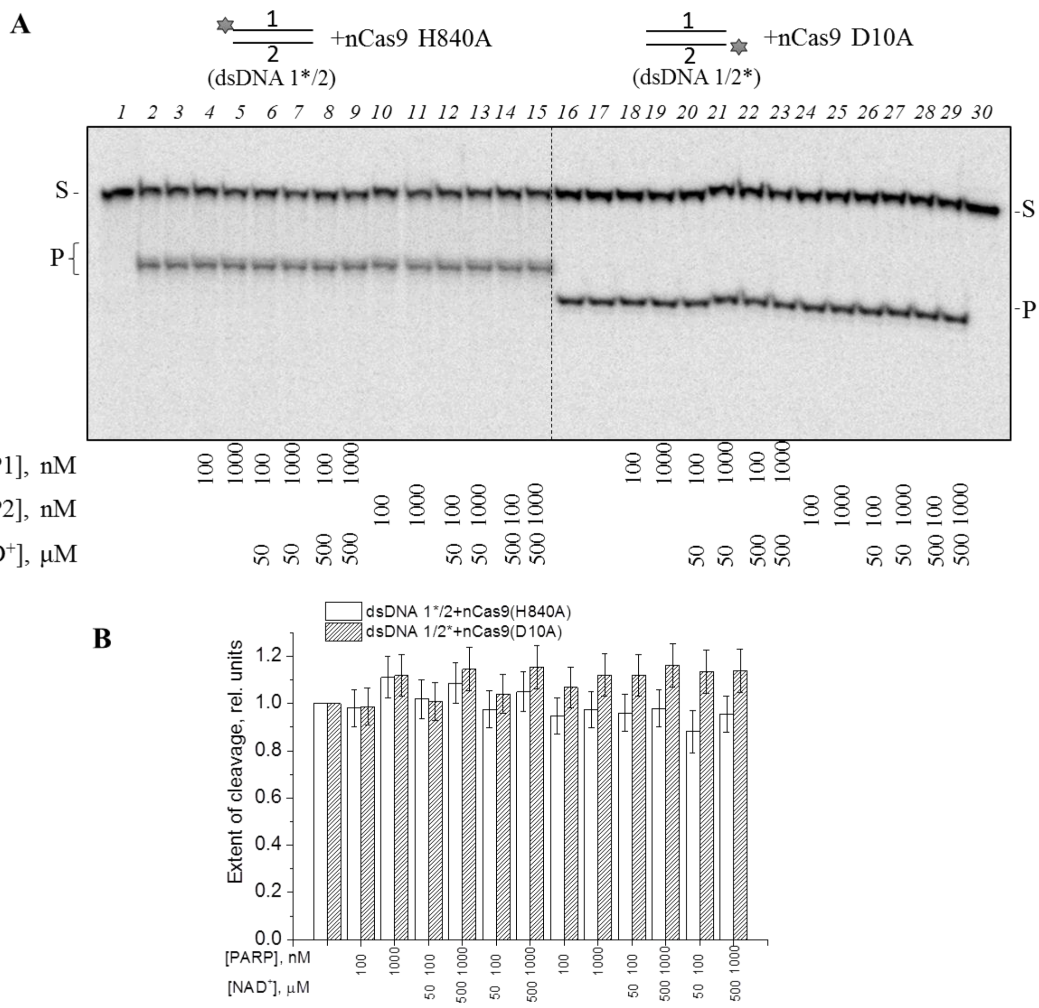


**S2 Fig. Cleavage of the oligonucleotide substrate by Cas9 nickase mutants in the presence of PARP1 and PARP2.** The endonuclease activity was tested by incubation of nCas9 H840A/sgRNA or nCas9 D10A/sdRNA (20 nM) with dsDNA 1*/2 or dsDNA 1/2* (10 nM) at 37°C for 30 min, in the absence (lanes 2, 3 and 16, 17) and presence (lanes 4–15 and 18–29) of PARP1/PARP2, without or with $NAD^+$. The reaction products were separated in a denaturing 20% PAG as shown in panel A. Bar charts (B) show the extent of dsDNA 1/2 cleavage induced by nCas9/sgRNA in the absence or presence of PARP1/PARP2 normalized to that in the absence of PARPs (the mean ± SD, n = 3). The 5’-$^{32}$P-labelled strand in dsDNA is asterisked.

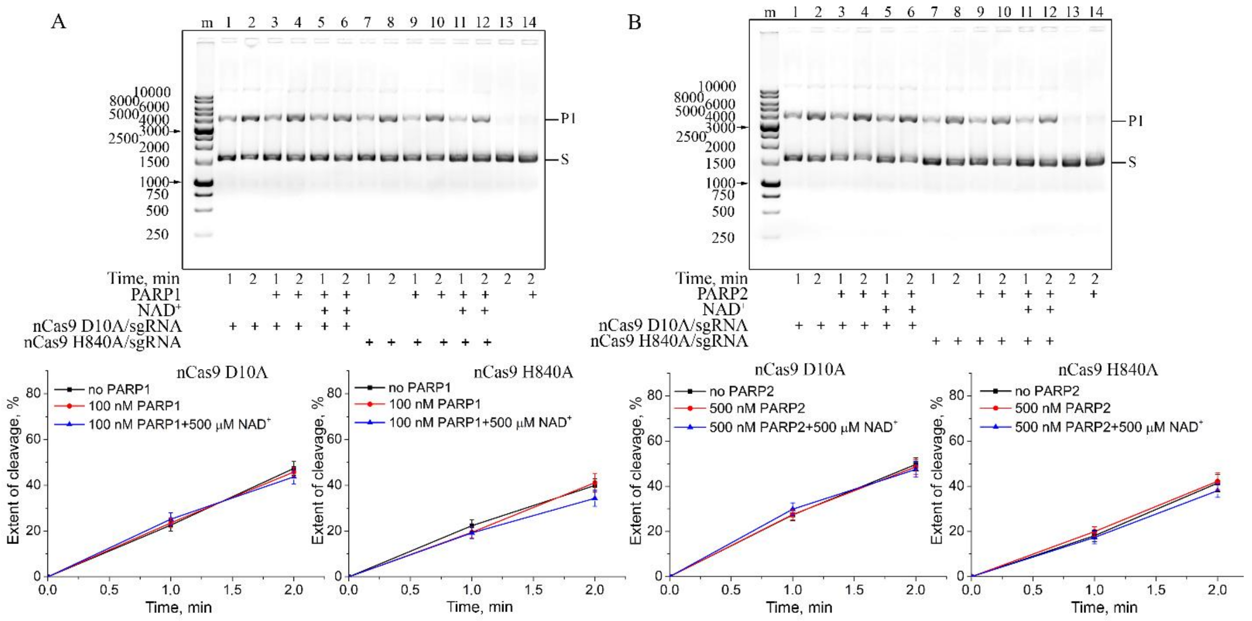

**S3 Fig. Cleavage of the plasmid substrate by Cas9 nickase mutants in the presence of PARP1 and PARP2.** The nCas9 D10A/sgRNA or nCas9 H840A/sgRNA complex (10 nM) was incubated with pLK1 DNA (10 ng/µl) at 37°C, in the absence (lanes 1, 2 and 7, 8) and presence (lanes 3–6 and 9–12) of PARP1 (100 nM) or PARP2 (500 nM) supplemented with 500 µM NAD$^+$ or not. The single-strand cleavage product (P1) was separated from the substrate (S) by electrophoresis in 1% GelRed stained agarose gel. The plots show the accumulation of cleavage product under the indicated conditions (the mean ± SD, n = 3).

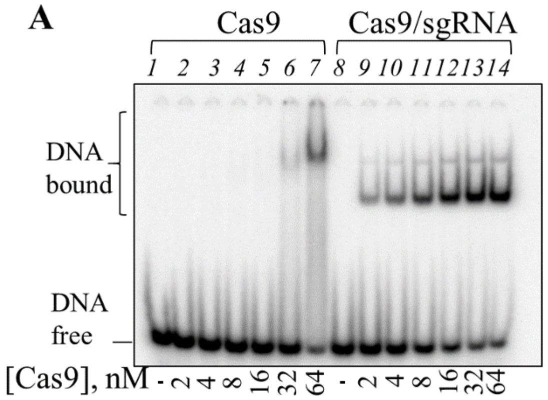


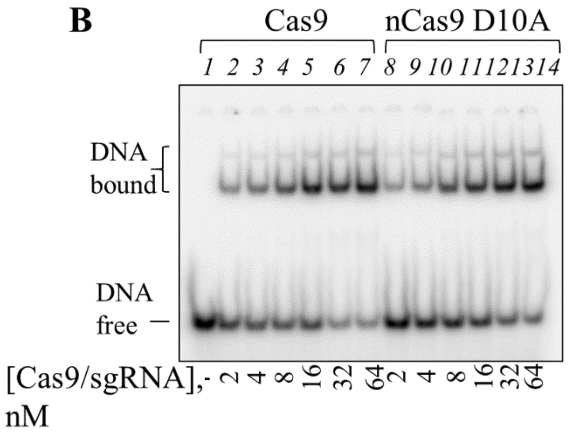


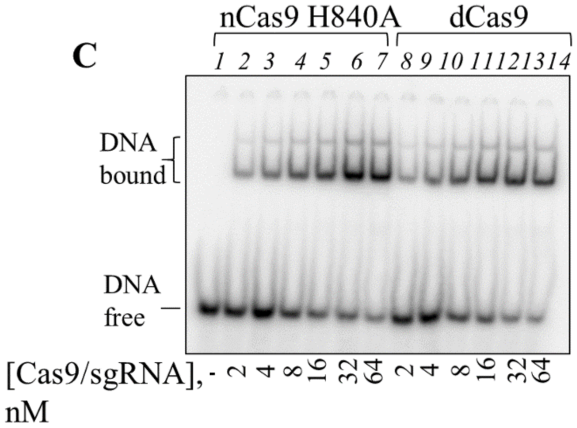


**D**

| Cas9/sgRNA | $K_d$, nM |
|---|---|
| Cas9 | 7.7±0.4 |
| nCas9 D10A | 9±1 |
| nCas9 H840A | 9±1 |
| dCas9 | 7±1 |

**S4 Fig. Binding of Cas9 and its mutant forms to the dsDNA substrate of Cas9**. The EMSA analysis of dsDNA 1/2* binding to Cas9 and Cas9/sgRNA (A), Cas9/sgRNA and nCas9 D10A/sgRNA (B), nCas9 H840A/sgRNA and dCas9/sgRNA (C). The reaction mixtures containing 10 nM dsDNA 1/2* and Cas9/Cas9 mutant (free or in the complex with sgRNA) at varied concentrations were incubated in the absence of $Mg^{2+}$ at 4°C for 30 min and separated in a native 5% PAG. The apparent $K_d$ values of the complexes (the mean ± SD, *n* = 3) are presented in Table (D).

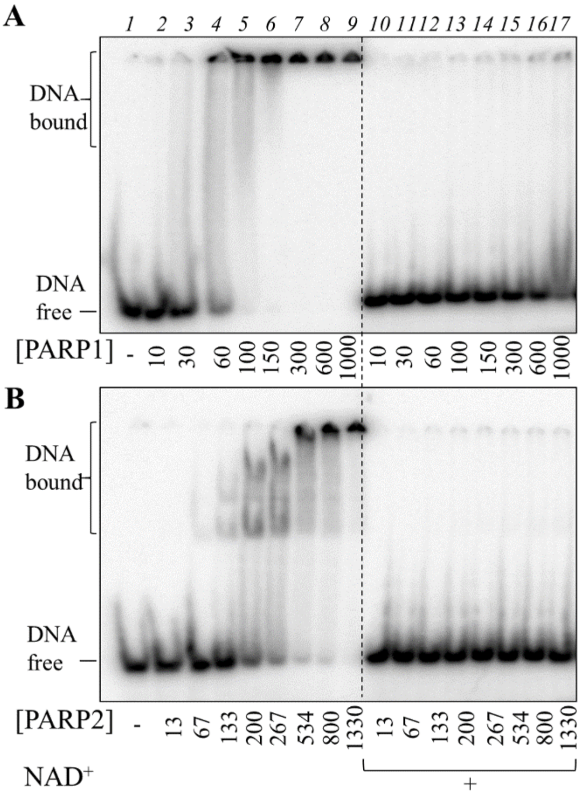


**C**

| PARP | $K_d$, nM |
|---|---|
| PARP1 | 53±5 |
| PARP2 | 200±16 |

**S5 Fig. Binding of PARP1 and PARP2 to the dsDNA substrate of Cas9**. The reaction mixtures containing 10 nM dsDNA 1/2* and PARP1 (A)/PARP2 (B) at the indicated concentrations were incubated in the absence (lanes 1–9) or presence (lanes 10–17) of 500 μM $NAD^+$ at 4°C for 30 min (in the absence of $Mg^{2+}$) and separated in a native 5% PAG. The apparent $K_d$ values of the complexes (the mean ± SD, *n* = 3) are presented in Table (C).

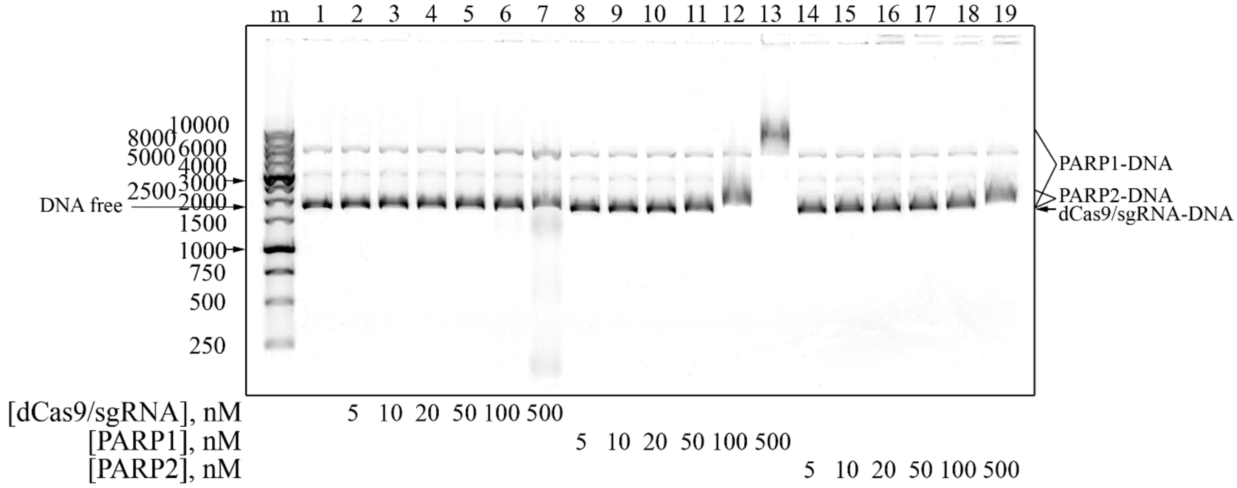


**S6 Fig. Binding of dCas9/sgRNA complex, PARP1 and PARP2 to plasmid DNA**. The pLK1 plasmid DNA (10 ng/µl) was incubated in the absence and presence of increasing concentrations of dCas9/sgRNA, PARP1 or PARP2. EMSA for the protein-DNA complexes was performed by electrophoresis in 1% GelRed stained agarose gel.

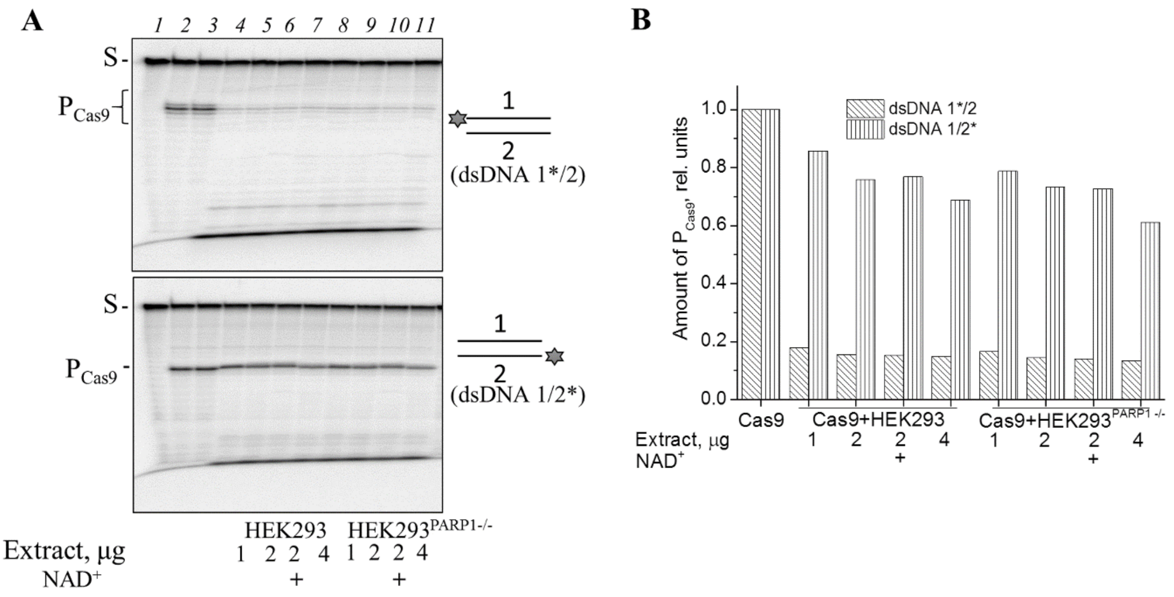


**S7 Fig. Protection of Cas9-generated dsDNA1/2 cleavage product from degradation in cell extracts.** Cas9/sgRNA (10 nM) was incubated with dsDNA1*/2 or dsDNA1/2* (10 nM), HEK293 or HEK293 *PARP1*$^{-/-}$ cell extracts and 500 μM NAD$^{+}$ as indicated. Cas9/sgRNA was pre-incubated with the substrate on ice for 60 min without $Mg^{2+}$, then the reaction mixtures were supplemented with cell extracts and 10 mM $MgCl_2$ and further incubated at 37°C for 30 min. (A) Representative gel images. (B) Relative yield of Cas9-cleaved DNA ($P_{Cas9}$) in the presence of cell extracts normalized to that in the absence of the extracts.

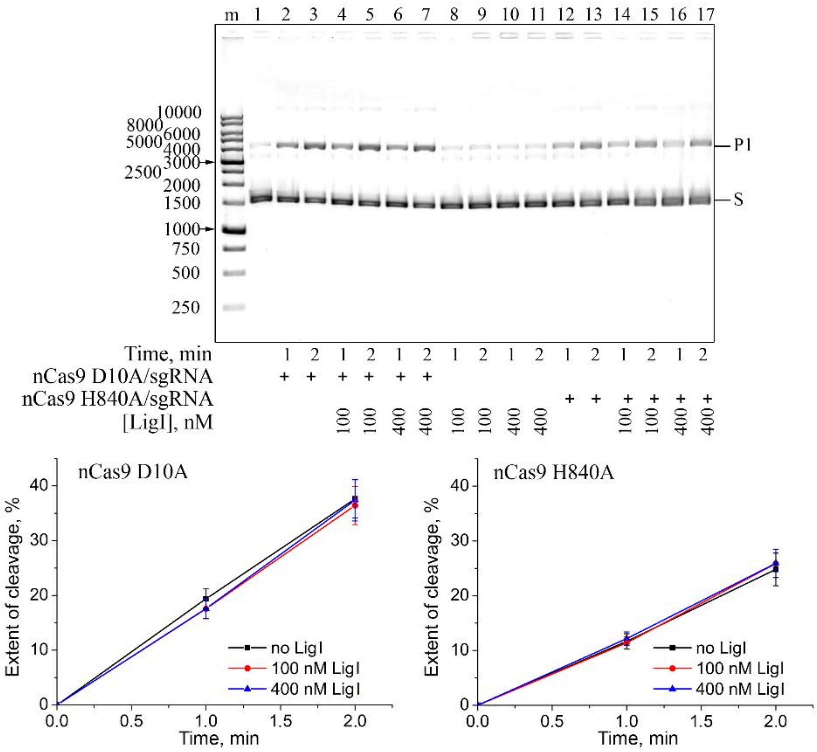


**S8 Fig. Effects of DNA ligase I on the nickase activity of Cas9.** (A) nCas9 D10A/sgRNA (10 nM) or nCas9 H840A/sgRNA (10 nM) was incubated with pLK1 DNA (10 ng/µl) at 37°C, in the absence and presence (+) of LigI (100 or 400 nM) (as detailed in Methods). The product (P1) was separated from the substrate (S) by electrophoresis in 1% GelRed stained agarose gel. The plots show accumulation of the cleavage product in the absence and presence of LigI (the mean ± SD, n = 3).

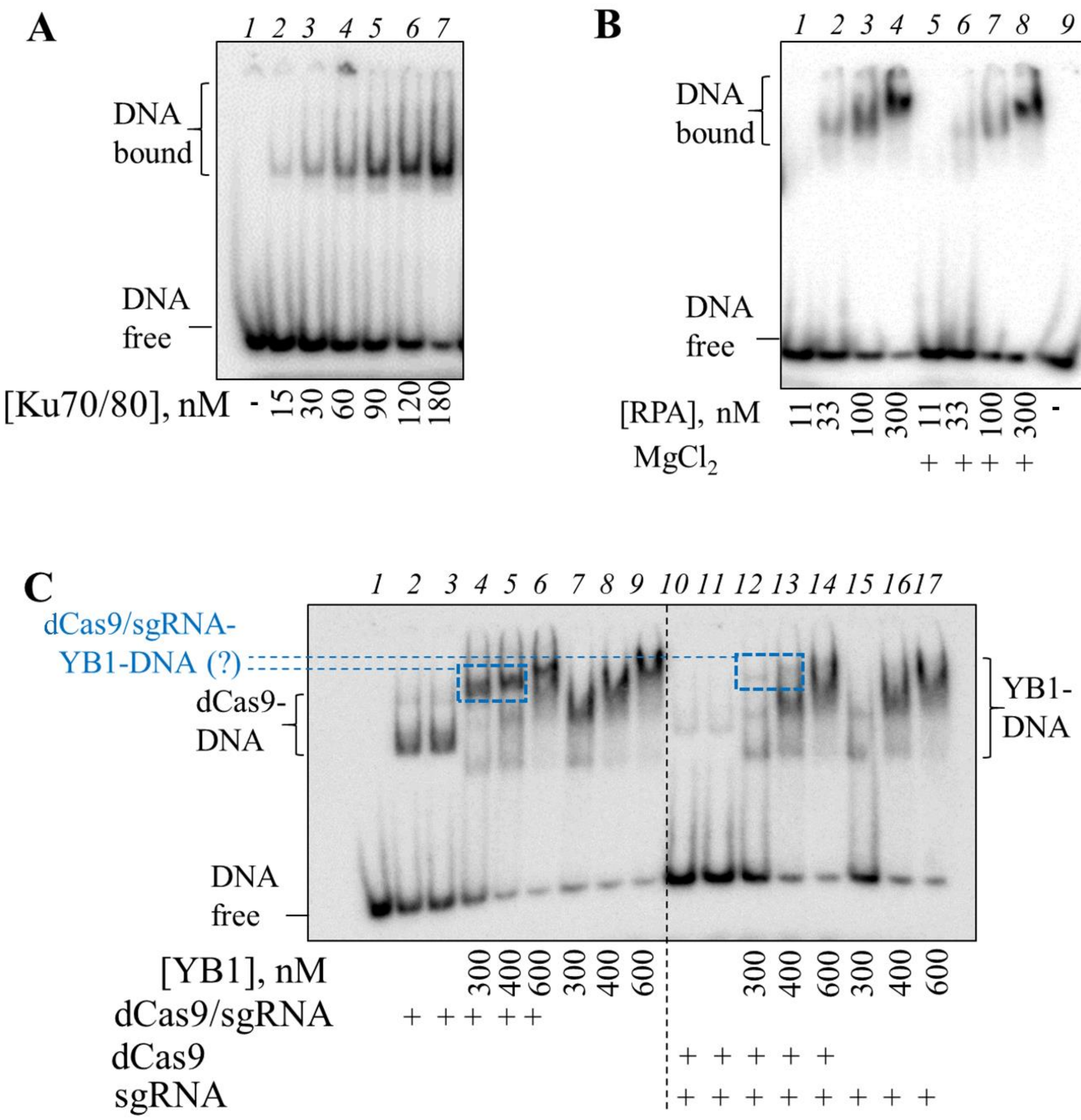


**S9 Fig. Binding of Ku70/80, RPA and YB1 to the dsDNA substrate of Cas9.** The reaction mixtures containing 10 nM dsDNA 1/2* and Ku70/80 (A) or RPA (B) at increasing concentrations were incubated at 4°C for 30 min and separated in a native 5% PAG. Binding of RPA was explored in the absence or presence of 10 mM $MgCl_2$. The $K_d$ values for RPA were 90 ± 20 nM (without $Mg^{2+}$) and 125 ± 40 nM (with $Mg^{2+}$), respectively. The DNA binding activity of YB1 (C) was explored in the absence (lanes 7–9) and presence of dCas9 (20 nM) and sgRNA (20 nM), added separately or together.

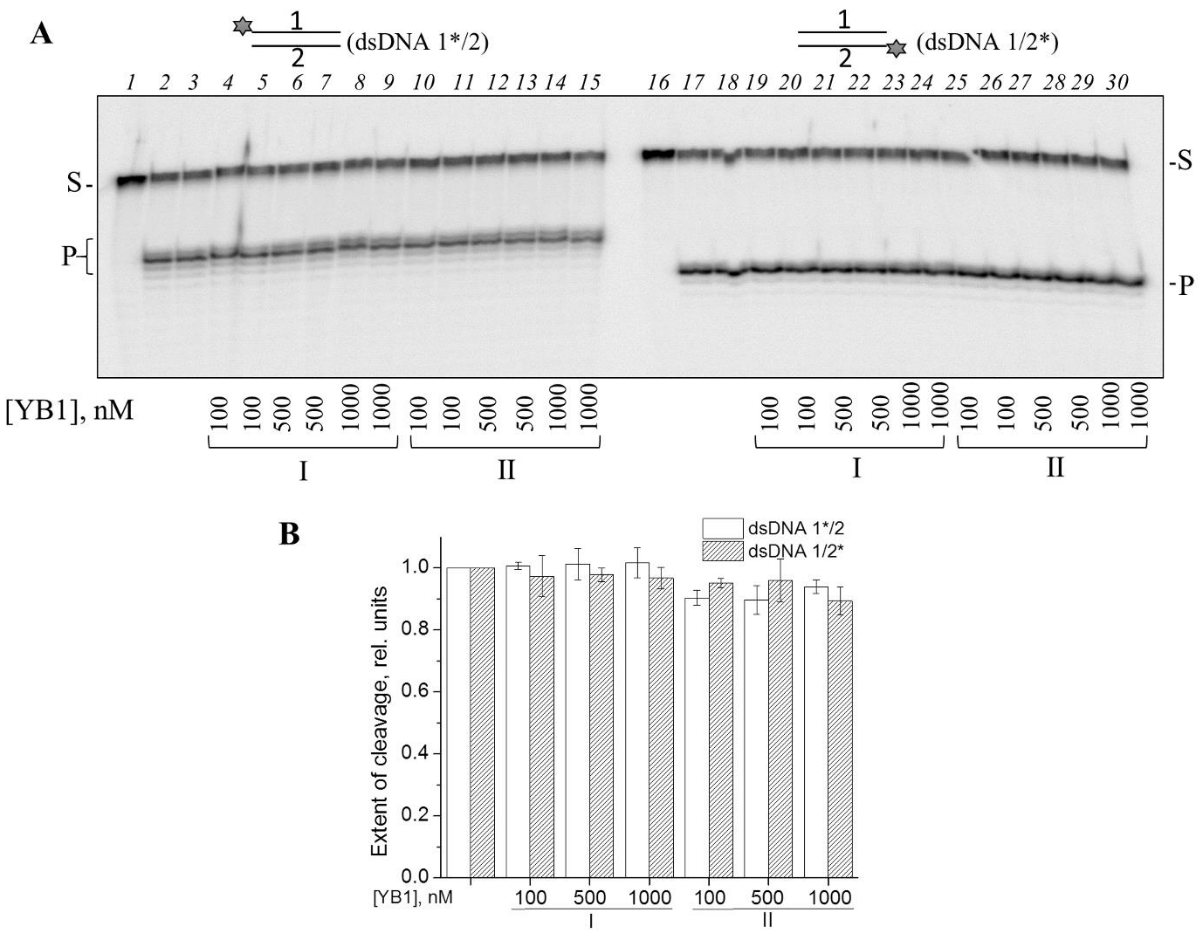


**S10 Fig. Effects of YB1 on the cleavage activity of Cas9**. (A) The endonuclease activity was tested by incubation of Cas9/sgRNA (20 nM) with dsDNA 1*/2 or dsDNA 1/2* (10 nM) at 37°C for 30 min, in the absence (lanes 2, 3 and 17, 18) and presence (lanes 4–15 and 19–30) of YB1 (100-1000 nM). In the presence of YB1, the reaction was performed without pre-incubation (I) or with pre-incubation of Cas9/sgRNA with YB1 on ice for 30 min and following addition of dsDNA 1/2 (II). The reaction products were separated in a denaturing 20% PAG. (B) Bar charts show the relative extent of dsDNA 1/2 cleavage induced by Cas9 in the specified reaction conditions (normalized to that in the control sample containing no YB1; the mean ± SD, *n* = 3).
